# a2iHelper: a Python toolkit for a differential editing site analysis of RNA-Seq data

**DOI:** 10.1101/2024.10.15.618547

**Authors:** Guilherme T. Ribas, Dieval Guizelini, Leonardo V. Riella, Cristian V. Riella

**Affiliations:** Center for Transplantation Science, Massachusetts General Hospital/Harvard Medical School, 149 13th st, Boston, 02129, MA, USA; Professional and Technological Education Sector, Federal University of Paraná, R. Dr. Alcides Vieira Arcoverde, 1225, Curitiba, 81520-260, PR, Brazil; Nephrology Division, Department of Medicine, Beth Israel Deaconess Medical Center, 330 Brookline Ave, Boston, 02215, MA, USA

**Keywords:** RNA editing, differential editing, *ADAR*

## Abstract

**Background:** A-to-I RNA editing, mediated by *ADAR* enzymes, plays a crucial role in cancer and autoimmune disorders but lacks standardized tools for differential analysis. After reviewing 55 studies, it highlights significant methodological heterogeneity, hindering result comparability and reproducibility. To address this, we developed a2iHelper, a Python package that streamlines RNA editing analysis by filtering noise, performing statistical analyses, and generating visualizations. a2iHelper integrates seamlessly with Python machine learning tools, aiming to standardize and simplify RNA editing research.

**Results:** We applied our methods to analyze A-to-I editing in a public dataset comparing wild-type and *ADAR* knockout. The source code is open, freely available on GitHub, and organized in a well-documented Python package. Using Snakemake for preprocessing, we conducted differential editing analysis on the top 104 most edited genes. The results include p-values from statistical tests, Odds ratios for Manhattan plots, and correlations between editing frequencies and gene expression, visualized in various plots.

**Conclusions:** We developed a2iHelper, a Python-based package for analyzing and visualizing A-to-I RNA editing data. It allows researchers with minimal programming experience to perform organized, reproducible editing analyses. Novice users can easily detect editing sites, filter noise, and generate figures, while advanced users can integrate functionalities into their workflows. a2iHelper runs on personal computers without needing High-Performance Computing resources.

## 1 Background

A-to-I RNA editing is a post-transcriptional modification in RNA transcripts where an Adenine (A) is replaced with an Inosine (I) by the action of the enzyme adenosine deaminase acting on RNA (*ADAR*) [1]. The editing process is associated with cancer and autoimmune disorders [2]. This substitution can occur in coding regions, leading to novel proteins by changing the codon to non-synonymous or inducing an alternative splicing. *ADAR* can also target non-coding regions, inducing instabilities in gene regulation, such as miRNA binding in 3’UTR regions. Also, the A-to-I substitution can prevent the immune response of double-strain RNA [1].

There are several tools to detect and count the A-to-I editing events using different strategies and focusing on distinct regions of the genome, such as REDItools2 [3], RnaEditingIndexer [4], SPRINT [5] and JACUSA2 [6]. However, there needs to be more tools dedicated to performing the differential editing analysis of this type of data in an easy scripting way. We reviewed the methodology of 55 research papers that performed A-to-I analysis, and we found a considerable heterogeneity between all steps of editing analysis approaches in different research groups (Fig A1 and Table A1). This heterogeneity doesn’t allow the community to compare the results from other groups, making it very hard to reproduce.

Therefore, we developed a2iHelper to assist researchers in running their workflow in a simple scripting language. Our package has several functions that can help detect editing sites, filter noise sites, perform statistical analysis, and generate figures. The entire package is based on the implementation in Python 3.10.13 and uses Pandas 1.5.3 [7] and NumPy 1.26.4 [8] libraries for data analysis. It makes it easy to integrate Python machine learning packages, such as Scikit-learning, for analysis.

## 2 Implementation

a2iHelper is divided into four principal modules: call reditools2, filter, stats, and plot. The call reditools2 provides an easy way to call REDI-tools2 [3] from the command line for different genome regions of interest. It also has methods to return editing from specific coordinates, such as 3’ UTR and 5’ UTR. The filter module merges output files from REDItools2, filtering noise positions such as low quality, low reads, missing values, and known SNPs.The stats provide methods to apply the most common statistical tests for analyzing differential data. The plot is a module dedicated to generating graphs with statistical test information when a test is performed. Fig 1 shows how the workflow is structured using a2iHelper modules. The package can be easily installed and freely available on GitHub (https://github.com/guilhermetabordaribas/a2iHelperPy).

**Fig. 1.**
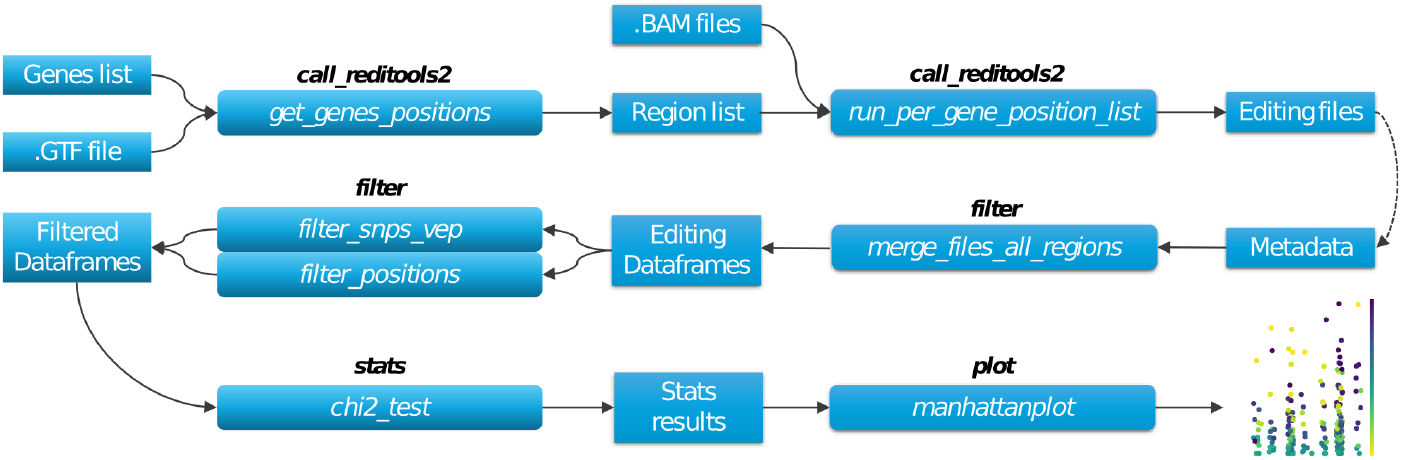
Workflow of the a2iHelper process. The terms in black are what the modules are called, and their functions are described inside the boxes. The result plot in this example is a Manhattan plot.

### 2.1 The call reditools2 module

This module enables running REDItools2 directly from your Python code or notebook. One needs to inform function the list of coordinates where in the genome the REDItools2 must analyze the editing, the complete path of the bam (Binary Alignment Map) file to be analyzed, the directory to save the editing files, the path of the reference genome sequence used to map the reads with transcript sequence and alignment stored to bam format file, the path where reditools.py is installed, a string with optional arguments to set the REDItools2 and the number of parallel jobs.

The Genomic coordinates list must have the chromosome name and the start and end positions (e.g., [‘chr2:122147686-122153083’, ‘chr18:60803848-60812646’]). Additionally, we provide an auxiliary method that enables the generation of coordinate lists in the described format from a list of genes annotated in GTF format, commonly utilized by alignment applications that generate BAM files.

### 2.2 The filter module

After running REDItools2, it is necessary to merge files from different samples and/or regions, keeping the information on the number of Adenine and Guanine and the editing frequency. The filter module has a function to do that; the user needs to inform a simple Pandas DataFrame with metadata about each editing file, where the first column is the path to editing files, the second column the names of the samples, the third column the region (gene or coordinates) and a fourth column with conditions of each sample (e.g., treated or wild type). The merging process also requires information on a minimum number of quality reads that should be considered in each edited site. The merging process will return three DataFrames: the first has A-to-I editing frequency, the second the Adenine counts per position, and the last one the Guanine counts per position.

The filter module has several functions that allow users to filter known SNP positions using the VEP Ensemble API automatically [9]. Also, it can be used to filter noising information in positions with a specified number of missing values, zero, and 100 percent editing frequency, as well as filtering positions by G-test of independence to verify if there’s deviation from the expected proportions and significant variation among the replicates to test if it is possible to pool samples from the same condition to perform posterior statical test.

### 2.3 The stats module

This module is a collection of statistical tests to perform differential editing levels between different conditions. Most of the tests are based on the Scipy 1.12.0 [10] package and are adapted to be easily applied to the editing data structure, requiring minimal data management from the user side. Besides statistical tests like the Mann-Whitney-U test, t-Student test, ANOVA, Kruskal-Wallis, Chi2, and Exact Fisher test, the user can compare the Shannon entropy of each position considering different conditions or not.

This module comprises a set of statistical tests designed to assess differential editing levels across distinct conditions. Most of these tests are grounded in the Scipy package and have been tailored for straight-forward application to the editing data structure, necessitating minimal data manipulation on the user’s part. In addition to conventional statistical tests such as the Mann-Whitney-U test, t-Student test, ANOVA, Kruskal-Wallis, Chi2, and Exact Fisher test, we include the Shannon entropy analysis at each position. The Shannon entropy offers a metric that may indicate the stability or instability observed at the edited site.

### 2.4 The plot module

This module is based on Matplotlib 3.8.4 [11] and Seaborn 0.11.2 [12] packages, and the functions automatically prepare the data to plot box plot, violin plot, bar plot, and Manhattan plot of genomic positions of user interest. Besides these graphs, it is possible to plot the heatmap of the correlation between the expression levels of *ADAR*, for example, and the editing frequency per position. The user can visualize the data, reducing the dimensionality with PCA and TSNE from the scikit-learning 1.4.1 package [13], and UMAP from umap-learn 0.5.5 [14].

## 3 Results and discussion

### 3.1 Example application

To demonstrate the methods above, we applied them to analyze A-to-I editing from a public dataset (GSE145011 PRJNA605715) [15]. In this demo, we analyzed the editing in wild type and with *ADAR* knock-out (*ADAR*-/-). The corresponding source code is found in Jupyter Notebooks. Alternatively, the raw Python source code is available in supplementals. The preprocessing was performed with a Snakemake workflow, which is also available.

For example, we conducted a differential editing analysis on the top 104 most edited genes sourced from the study GSE145011. We extracted the coordinates from the GTF annotation file based on the gene names using the function get genes positions. Then, we used the coordinates to run REDItools2 on these genes using the function run per gene position list. The output format of the RNA editing file (.res) is listed in Table 1. If the file was generated via REDItools2, the file is in the correct format. If another editing detector was used, ensure that the columns file is in the same order and type as in Table 1 (Region, Position, Reference, Strand, Coverage-q30, BaseCount[A, C, G, T]).

**Table 1.** RNA editing format to be use as input for a2iHelper.

| Region | Position | Reference | Strand | Coverage-q30 | BaseCount[A,C,G,T] |
| --- | --- | --- | --- | --- | --- |
| chr19 | 55401670 | C | 2 | 3944 | [0, 1847, 0, 2097] |
| chr19 | 55396137 | A | 2 | 4 | [2, 0, 2, 0] |
| chr19 | 55389165 | A | 2 | 337 | [197, 1, 139, 0] |
| chr19 | 55387928 | A | 2 | 119 | [87, 32, 0, 0] |
| chr19 | 55396022 | T | 2 | 4 | [0, 1, 0, 3] |

To merge all REDItools2 output “.res” files, one can use the merge files all regions informing the metadata DataFrame with at least the information of editing file path, name of the sample, region, and condition, in this order. Table 2 lists an example of metadata DataFrame.

**Table 2.** Example of metadata format.

| File name | Sample | Region | Condition |
| --- | --- | --- | --- |
| /.../SRR11050913_chr20_3846799_3876123_RES.tsv | SRR11050913 | MAVS | wild type |
| /.../SRR11050914_chr20_3846799_3876123_RES.tsv | SRR11050914 | MAVS | wild type |
| /.../SRR11050915_chr20_3846799_3876123_RES.tsv | SRR11050915 | MAVS | *ADAR1*-/- |
| /.../SRR11050916_chr20_3846799_3876123_RES.tsv | SRR11050916 | MAVS | *ADAR1*-/- |

After merging files, it is possible to apply filters for known SNPs, missing values, 0% and 100% of editing per condition or in total samples. It is easy to perform statistical tests like the Mann-Whitney U, Fisher exact, and Chi-square tests to retrieve the p-values. Fig 2-AB shows how two tests can return different statistically relevant differential positions. The a2iHelperPy also allows easy calculation of the Odds ratio to plot a Manhattan plot to visualize the association of each condition and the proportion of bases Adenine and Guanine (Fig 2-C). The user can also evaluate the base of Shannon entropy of each condition (Fig 2-D) as well the correlations of editing frequencies between conditions (Fig 2-E) and correlations between editing frequencies and gene expression (Fig 2-F).

**Fig. 2.**
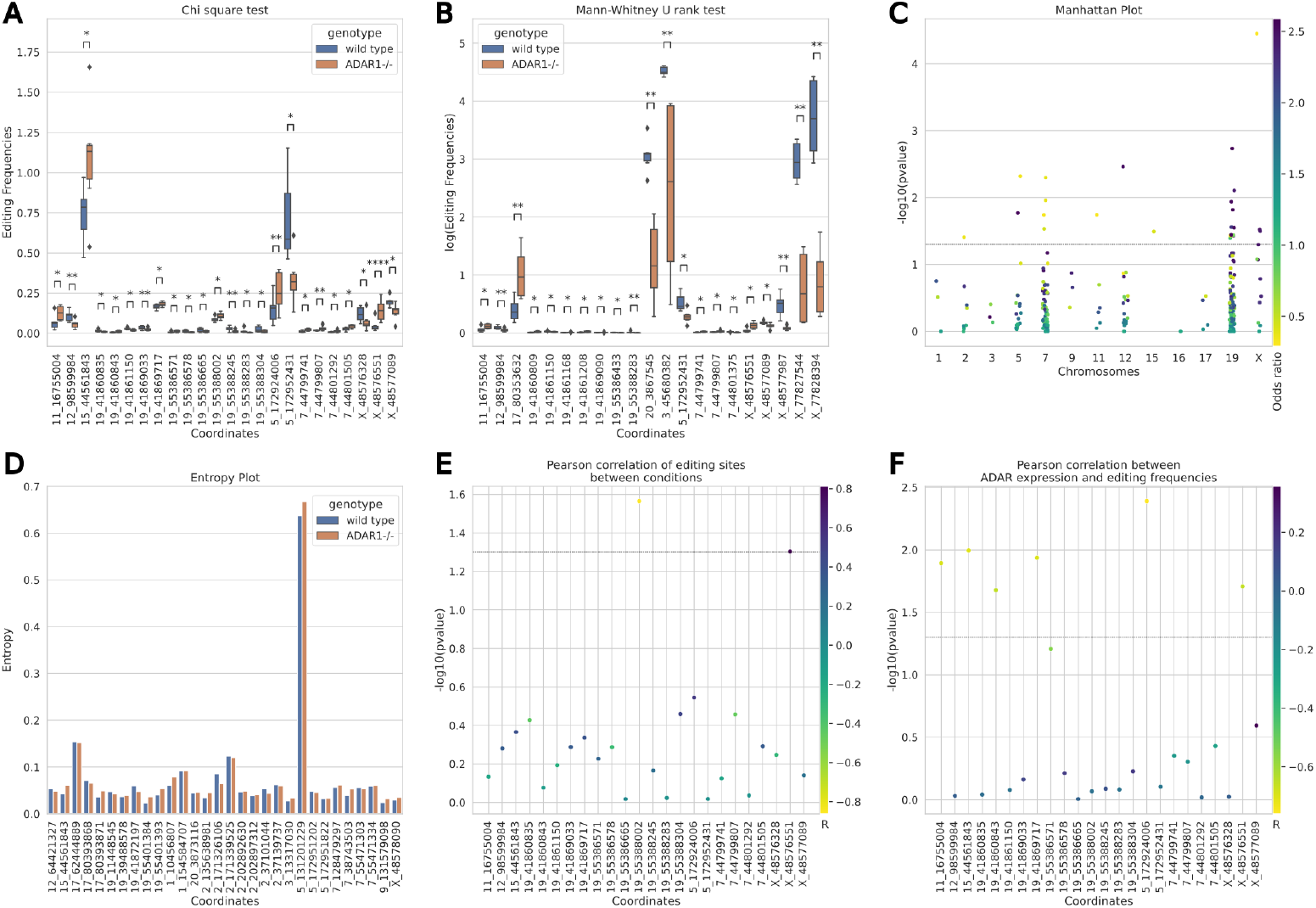
Example of resulting analysis for a2iHelper. (A) A boxplot of editing frequency distributions and the significance of the statistics are shown. A chi-square test between wild type and *ADAR*-/-was used to test the difference in the counts of Adenine and Inosine in each condition. (B) A boxplot of editing frequencies and statistical significance. A Mann-Whitney U rank test was used to test the difference in editing frequencies in wild type and *ADAR*-/-. (C) A Manhattan plot shows the Chromosome’s names on the horizontal axis and the -log10 of the p-value on the vertical axis. The color map describes the odds ratio of each sample. (D) A barplot representing the entropy of each analyzed coordinate and condition. (E) A scatterplot represents the coordinates in the horizontal axis, the vertical -log10 of p-value, and the color map of the value of the R Pearson correlation between editing frequencies in different conditions. (F) A scatterplot represents the coordinates on the horizontal axis, the vertical -log10 of the p-value, and the color map, which is the R Pearson correlation between editing frequencies and *ADAR* expression. The horizontal dotted line in C, E, and F represents the limit of a p-value *≤* 0.05. In A and B, the asterisks represent the significance of the p-value. The asterisks indicate statistical significance: * 0.01 *<* p *≤* 0.05, ** 0.001 *<* p *≤* 0.01, *** 0.0001 *<* p *≤* 0.001 and ****: p *≤* 0.00001.

## 4 Conclusion

We developed the a2iHelper to provide a Python-based package to perform statistics analysis and visualization of A-to-I RNA editing data. a2iHelper enables experimental biologists with little computational programming experience to conduct an editing analysis in a well-organized and reproducible way. Novice programmers can follow the available documentation and easily detect editing sites, filtering noise sites and SNPs, statistical analysis, and figures generation. Advanced users can integrate a2iHelper functionalities in their algorithms and skip writing code for general filtering sites’ differential statistics analysis. The entire workflow doesn’t require High Performance Computing (HPC) resources and can be run on a personal computer.

## 5 Availability and requirements

- Project name: a2iHelper
- Project home page: https://github.com/guilhermetabordaribas/a2iHelperPy/
- Package documentation: https://a2ihelperpy.readthedocs.io/
- Code of paper version: https://doi.org/10.7910/DVN/NUXBR9
- Code of example application: https://doi.org/10.7910/DVN/NP9MXF
- Operating system(s): Ubuntu 22.04.3 LTS
- Programming language: Python 3.10.13

## Declarations

The authors declare no competing interests.

## Appendix A Supplementary Figures and Tabels

**Fig. A1.**
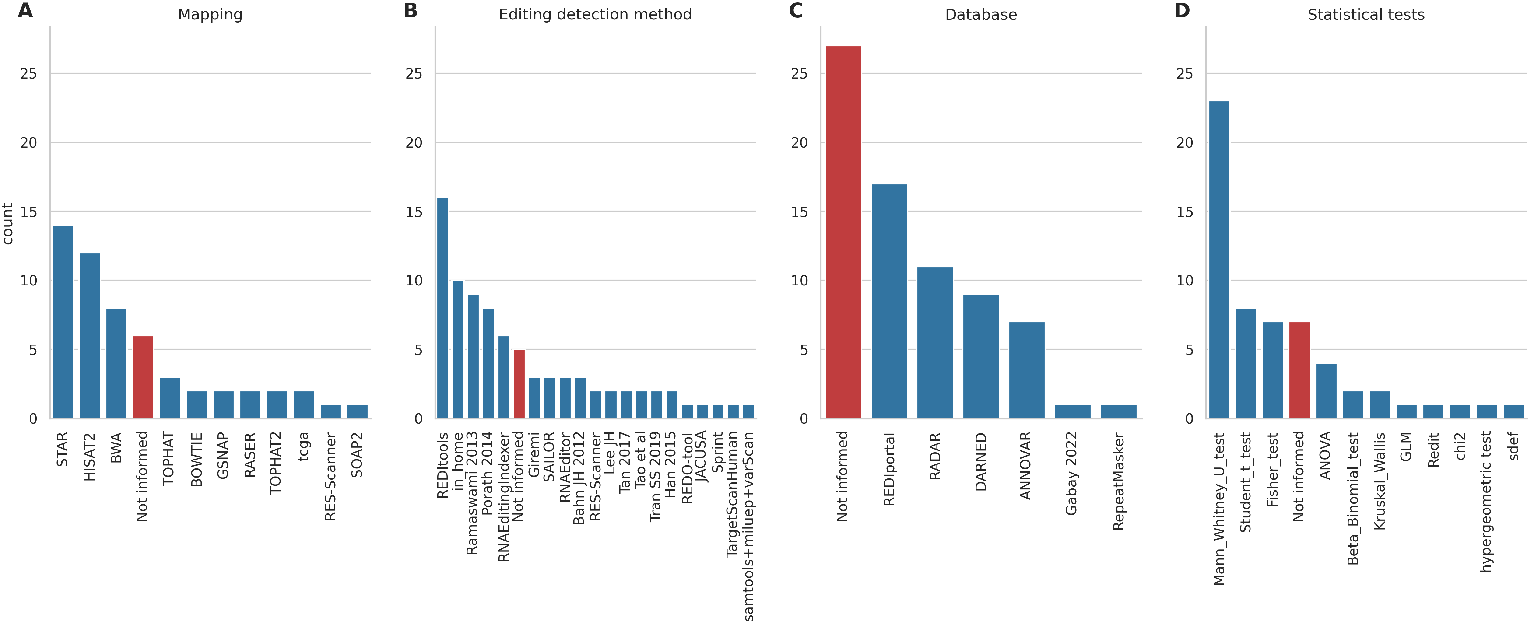
Workflow of the a2iHelper process. The terms in black are what the modules are called, and their functions are described inside the boxes. The result plot in this example is a Manhattan plot.

**Table A1.** Publications about editing in RNA editing (Figure A1)

| Title | Publication year | PubMed ID |
| --- | --- | --- |
| Genome-wide perturbations of Alu ex... | 2022 | 35042936 |
| RNA Editing in Glioma as a Sexually... | 2022 | 35406793 |
| Differential Analysis of A-to-I mRN... | 2022 | 35052353 |
| Preliminary identification and anal... | 2022 | 35342974 |
| Brain Epitranscriptomic Analysis Re... | 2022 | 35559016 |
| RNA Editing-Associated Post-Transcr... | 2022 | 35462874 |
| Genome-Wide Analysis of Multiple Or... | 2022 | 35050036 |
| A-to-I nonsynonymous RNA editing wa... | 2022 | 36064557 |
| Editome landscape of CCM-derived en... | 2022 | 35771000 |
| Repeated Winning and Losing Experie... | 2022 | 35664469 |
| A game changer for bipolar disorder... | 2022 | 35504874 |
| Characterization of RNA Editome in ... | 2022 | 35049829 |
| RNA editing signatures identify mel... | 2021 | 39225857 |
| Profiling neural editomes reveals a... | 2021 | 33355311 |
| RNA editing regulates lncRNA splici... | 2021 | 34851956 |
| Integrative analysis of long extrac... | 2021 | 33391469 |
| Long Term Maturation of Human Corti... | 2021 | 33619405 |
| RNA editing at a limited number of ... | 2021 | 33983932 |
| Quantifying RNA Editing in Deep Tra... | 2020 | 32211029 |
| Differential RNA Editing and Intron... | 2020 | 33317061 |
| HyperTRIBE uncovers increased MUSAS... | 2020 | 32332729 |
| Differential RNA editing between ep... | 2020 | 33106178 |
| RNA editing in cancer impacts mRNA ... | 2020 | 33106178 |
| ADAD1 and ADAD2, testis-specific ad... | 2020 | 32665638 |
| Dynamic inosinome profiles reveal n... | 2019 | 30760294 |
| Genomic positional dissection of RN... | 2019 | 30949194 |
| Regulation of RNA editing by RNA-bi... | 2019 | 30652130 |
| Adenosine-to-Inosine RNA Editing in... | 2019 | 31576663 |
| Genome-wide identification and anal... | 2019 | 31158229 |
| Widespread RNA editing dysregulatio... | 2019 | 30559470 |
| Genome-wide identification and anal... | 2018 | 29470549 |
| Increased RNA Editing May Provide a... | 2018 | 29617672 |
| Genome-wide analysis of differentia... | 2017 | 28250018 |
| A-to-I editing in human miRNAs is e... | 2017 | 28550310 |
| Regulation of gene expression and R... | 2017 | 29146998 |
| The C. elegans neural editome revea... | 2017 | 28925356 |
| Whole plastid transcriptomes reveal... | 2017 | 28916985 |
| RNA editing signature during myeloi... | 2017 | 28484266 |
| Testicular adenosine to inosine RNA... | 2017 | 28395340 |
| Dynamic landscape and regulation of... | 2017 | 29022589 |
| Structure-mediated modulation of mR... | 2017 | 29093448 |
| Adenosine-to-inosine RNA editing co... | 2016 | 27595325 |
| Complex regulation of ADAR-mediated... | 2016 | 26768488 |
| Dynamic regulation of RNA editing i... | 2016 | 27348216 |
| Genome-wide identification of RNA e... | 2015 | 25462863 |
| The Genomic Landscape and Clinical ... | 2015 | 26439496 |
| Profiling RNA editing in human tiss... | 2015 | 26449202 |
| Elevated RNA Editing Activity Is a ... | 2015 | 26440895 |
| Principles Governing A-to-I RNA Edi... | 2015 | 26440892 |
| Comparative RNA Editing in Autistic... | 2013 | 22869036 |
| Identifying RNA editing sites using... | 2013 | 23291724 |
| Recoding RNA editing of antizyme in... | 2013 | 23291631 |
| Accurate identification of human Al... | 2012 | 22484847 |
| Widespread RNA and DNA Sequence Dif... | 2011 | 21596952 |
| Alu Sequences in Undifferentiated H... | 2010 | 20574523 |

